# Alpha Neurofeedback Training with a portable Low-Priced and Commercially Available EEG Device Leads to Faster Alpha Enhancement: A Single-blind, Sham-feedback Controlled Study & Methodological Review

**DOI:** 10.1101/527598

**Authors:** Adrian Naas, João Rodrigues, Jan-Philipp Knirsch, Andreas Sonderegger

## Abstract

**Introduction:** Findings of recent studies have proposed that it is possible to enhance cognitive capacities of healthy individuals by means of individual upper alpha (around 10 to 13.5 Hz) neurofeedback training. Although these results are promising, most of this research was conducted based on high-priced EEG systems developed for clinical and research purposes only. This study addresses the question whether such effects can also be shown with an easy to use and comparably low priced Emotiv Epoc EEG headset available for the average consumer. In addition, critical voices were raised regarding the control group designs of studies addressing the link between neurofeedback training and cognitive performance. Based on an extensive literature review revealing considerable methodological issues in an important part of the existing research, the present study addressed the question whether individual upper alpha neurofeedback has a positive effect on alpha amplitudes (i.e. increases alpha amplitudes) and short-term memory performance focussing on a methodologically sound, single-blinded, sham controlled design.

**Method:** Participants (N = 33) took part in four test sessions over four consecutive days of either neurofeedback training or sham feedback (control group). In the experimental group, five three-minute periods of visual neurofeedback training were administered each day whereas in the control group, the same amount of sham feedback was presented. Performance on eight digit-span tests as well as participants’ affective states were assessed before and after each of the daily training sessions.

**Results:** Participants in the neurofeedback training (NFT) group showed faster and greater alpha enhancement compared to the control group. Contrary to the authors’ expectations, alpha enhancement was also observed in the control group. Surprisingly, exploratory analyses showed a significant correlation between the initial alpha level and the alpha improvement during the course of the study. This finding suggests that participants with high initial alpha levels profit more from alpha NFT interventions. digit-span performance increased in both groups over the course of time. However, the increase in individual upper relative alpha did not explain significant variance of digit-span improvement. In the discussion, the authors explore the appearance of the alpha enhancement in the control group and possible reasons for the absence of a connection between NFT and short-term memory.

## 1. Introduction

An alarming indicator for the need of cognitive enhancement in our society is the growing number of college students using drugs like Methylphenidate (MPH, Ritalin) or Modafinil to enhance concentration, memory performance and wakefulness (16% on some college campuses, see e.g. [1–3]). Rather than demonising the need for cognitive enhancement, the aim of this piece of research is to examine the usefulness of the non-invasive technique of neurofeedback training (NFT) for cognitive enhancement. Previous research addressing this question has reported some evidence for a positive relationship. However, to the authors’ knowledge, no study has tested the effectiveness of NFT with a not-medical grade EEG. This gap in current research, leaves the average consumer torn between the glorious slogans of a booming BCI industry with their easy-to-use and low-priced devices and a common sense which tries to disentangle advertising from possibility and innovation. On top of that, the task is complicated by science, which works beyond the omnipresent publication bias [e.g. 4] with methodologically problematic designs like no-intervention control groups.

Summing up, this piece of research focuses on two aspects. Firstly, it aims at the investigation of the effectiveness of alpha NFT with an easy to use and low priced EEG Headset and a corresponding software. Secondly, it provides an overview regarding methodological aspects in the field of alpha NFT and cognitive enhancement. Or, to put it differently, the authors humbly try to support average consumers when they are faced with questions like “If I train with an *Emotiv Epoc* and the corresponding software, will my short-term memory get better?”.

But before we can answer this question, a short introduction into NFT shall be given. NFT is a process during which subjects learn to influence their EEG pattern, for example by enhancing their individual upper alpha (IUA) amplitude [5]. However, other EEG components like the amplitude of different EEG frequency bands e.g. theta, alpha or beta can be fed back as well. The feedback can be provided as bar graph [6], colour code [7] or as a function of different sounds [8]. In combination with a mental strategy (e.g. thinking about friends, [9]), the users can shape their brain activity into a certain direction (for instance enhance alpha activity), which in the case of alpha is considered to be beneficial for cognitive performance [10]. NFT can be seen as non-invasive technique to alter brain activity. Unlike for example transcranial magnetic stimulation (TMS), NFT does not interfere actively with the brain but serves merely as a mirror of the current amplitude of the target frequency band.

To sum up, NFT can be described as a process during which neural activity is consecutively shaped into a predefined direction, by applying a mental strategy (e.g. visualizing engaging in a hobby) which is adjusted during a circular learning process (see Figure 3), based on a EEG feedback (e.g. colour scheme, sounds), in combination with conditioning procedures (reward processes, symbols or sounds). This process of altering oscillatory cerebral activity to increase the individual upper alpha amplitudes can, according to some authors [5,11,12], positively influence cognitive performance.

For a contribution to the understanding of the connection between NFT and cognitive enhancement, the study at hand follows three different purposes. Firstly, we aimed at replicating findings of previous experiments which indicated beneficial effects of individual upper alpha NFT on cognitive performance [5,13], while secondly taking a step into the direction of the average consumer by choosing an affordable device which is easy to use. Thirdly, an overview of methodological aspects like control group designs and blindfolding of publications in the field of alpha NFT and cognitive performance shall be given. Before these three areas of interest are explored more in detail, a short introduction into the origin of NFT is be given.

### 1.1. The Origin of Neurofeedback Training

The use of NFT in medical and therapeutic contexts has gained increasing interest in research and practice over the past 50 years (e.g. [14–17]). Various studies indicated that NFT shows positive effects in the treatment of diseases like Attention Deficit Hyperactive Disorder, Autism Spectrum Disorder, Substance Use Disorder and Epilepsy (e.g. [14–24]). Also with regard to other disorders (e.g. General Anxiety Disorder, see [29]), there are some studies suggesting positive effects of NFT. Recently, a first pilot study investigating the usefulness of NFT as intervention technique for patients suffering from Alzheimer Disease (AD) revealed that “neurofeedback, in combination with treatment with cholinesterase inhibitors, may be a potential treatment by which the progressive deterioration in patients with AD can be stabilized” [30]. Finally, NFT seems to facilitate effectively the lives of people affected by Amyotrophic Lateral Sclerosis or the so-called Locked-in Syndrome [31].

In the field of therapeutic application of NFT, one of the best established application domains is the treatment of Attention Deficit Hyperactive Disorder. Different studies have shown positive results of a NFT intervention. For example Linden et al. [32] conducted a study with an intensive training schedule of forty 45-min sessions of NFT over a period of 6 months. The training aimed at enhancing beta activity and supressing theta activity at the electrode sites Cz and Pz (international 10-20 system). NFT was given by means of computer games and conditioning was enhanced by rewarding the participants with small gifts after the intervention, if the level of performance was satisfying. After the training course, the experimental group performed significantly better compared to the waiting list control group and compared to the individual pre-course measurement on both, an IQ-Test (*K-Bit IQ*) and a parent behaviour rating scale for inattention.

### 1.2. Alpha Neurofeedback Training for Cognitive Enhancement in Healthy Participants

Because of its positive effects in clinical practice, there has been increasing interest in the question whether NFT can influence the capacities of healthy individuals positively as well. Some studies seem to support this hypothesis [33] and according to Klimesch [12], especially the individual upper alpha band is of major importance for cognitive performance.

The individual upper alpha band is generally calculated based on EEG data. By means of Fast Fourier Transformation (FFT) the rhythmic EEG components delta (about 0.5 – 4 Hz), theta (about 4 – 8 Hz), alpha (about 8 – 13 Hz) and beta (about 13 – 30 Hz) can be extracted. The IUA constitutes a sub-band of the alpha component and is located between the individual alpha peak (IAP, between 7.5 and 12.5 Hz) and IAP + 2 Hz [12].

The ‘amount’ of alpha activity can be expressed in terms of amplitude or power. Working with amplitude instead of power values has the advantage that it prevents excessive skewing and improves the validity of the statistical analysis [6]. Sometimes (e.g. [34]), instead of amplitude values, relative alpha values are calculated by dividing the mean amplitude of the individual upper alpha band by the mean amplitude of the whole EEG. This normalization avoids variance in the absolute alpha amplitude caused by changes between trials due to changes in impedance between the electrodes and the scalp. By normalizing, the frequency band of interest is relativized, which mitigates the issue of attenuations caused by external factors, which affect all frequency bands equally [6].

Alpha is an especially interesting oscillation that the human brain exhibits. It’s the predominant rhythm in the human brain in a resting state, especially when eyes are closed [35]. Until the 80’s, Alpha NFT was considered as a simple relaxation training, located within the theoretical framework of unitary arousal models. Only during the 90’s, new interest arose and from then on many different research questions circulated around the alpha frequency band [36], which will be outlined in the following paragraphs.

The first alpha property we want to consider is the association between individual alpha peak position and cognitive performance and neurological disorders, respectively. After conducting a FFT, the data can be plotted in a frequency spectrum map. In resting state recordings, the alpha peak is clearly visible between 7.5 and 12.5 Hz and constitutes one of the strongest components of the FFT. Higher alpha peak frequencies (e.g. 12 Hz in comparison to 10 Hz) correlate negatively with neurological disorders and with low age and high age. Furthermore, higher alpha peak frequencies correlate positively with high memory performance [37,38] and IQ [39].

Another property of the alpha activity is the connection between alpha amplitude/power and cognitive performance. For example Neubauer and colleagues [40] found a positive correlation between individual upper alpha amplitude and IQ. More specifically, high alpha power during a resting state and low alpha power during the execution of a task was associated with good performance in semantic longterm memory tasks [12]. According to Klimesch [12], alpha shows a task-related desynchronization, it increases during resting states (especially when eyes are closed) and decreases during performance of a cognitive task (e.g. mental calculations). Therefore, it seems to be a promising approach to mimic the phenomena observed in good performers by means of NFT (enhanced alpha power during a resting period shortly before the short-term memory task) in order to enhance cognitive performance.

Interestingly, past studies [12] observed the connection between alpha desynchronization and cognitive performance only when the alpha band was divided into two sub-bands: upper and lower alpha. Klimesch located the upper alpha band between the individual alpha peak (IAP, between 7.5 and 12.5 Hz) and IAP + 2 Hz and stated that the lower alpha band is connected to a “variety of non-task and non-stimulus specific factors which may be subsumed under the term ‘attention’ […] and reflect general task demands” [12]. This author located the lower alpha band between IAP - 4 and IAP. Therefore, in most of the studies, the individual upper alpha (IUA) band was used for NFT and some researchers go as far as assessing the alpha band each test session anew.

A study of high importance for the development of IUA feedback addressed the topic by means of transcranial magnetic stimulation in a within-subject design [33]. In line with the correlational findings between alpha desynchronization and cognitive performance, the participants were stimulated to produce more IUA activity (individual alpha peak + 1 Hz) at P6 and Fz before the execution of a task. In this way, the natural desynchronization process which can be found in participants showing high cognitive performance (i.e. mental rotation and short-term memory performance) was mimicked. In the control condition, participants also ‘underwent’ transcranial magnetic stimulation, but the coil was rotated by 90° so the participants did not receive any stimulation. The results show a significant increase of IUA during transcranial magnetic stimulation in the experimental condition, as well as decreased test power, resulting in a large event-related desynchronization. None of these changes were observed in the control condition. Cognitive performance was assessed in terms of success in a *Mental Rotation Task.* Results showed that mental rotation performance in the experimental condition was higher compared to the control condition. The authors interpreted these findings as an indicator for a causal relationship between IUA activity and cognitive performance in healthy subjects.

Based on these findings, several studies examined the connection between IUA activity and cognitive performance. In those studies, different aspects of cognitive performance like short-term memory performance_or working memory performance were assessed via a *digit-span Task* or a *Conceptual Span Task* (e.g. [5,41]), or *Mental Rotation Task* [42]. Mental flexibility and executive functions were assessed via the *Trail Making Test* [43], or creativity by the *Unusual Uses Test* [44]. Summarizing the results of these studies, imply a positive connection between individual upper alpha NFT and different aspects of cognitive performance like working memory/STM and visuospatial rotation. Whether the relationship between IUA and STM is of causal or correlational character, which underlying mechanisms lead to the enhancing effect of individual upper alpha NFT on cognitive performance and whether unspecific environmental factors of the experimental setup play a key role in the process of NFT is still not fully understood at the moment. In the following section, some of these aspects are addressed by a comprehensive analysis of published studies addressing the link between IUA and cognitive performance.

### 1.3. Summary of Experimental Studies on Neurofeedback Training and Cognitive Performance

This section summarizes findings of studies addressing the link between IUA and cognitive performance. Inspired by Rogala and colleagues [45], Table 1 gives an overview of the existing experimental research addressing NFT training (Alpha and Alpha/Theta) and its effects on behavioural measurements of attention (column “A”) and memory (column “M”). Column “G” represents general success and subsumes general effects of the training obtained in any of the investigated measures other than memory and attention. Studies regarding Alpha NFT and Memory were highlighted grey and methodological aspects which deserve critical attention are marked bold. The overview contains studies that appear in [45] (marked with an asterisk *) as well as new research that has not been considered in their review. Inclusion criteria were that the studies used alpha as feedback frequency and the dependent variable was not a clinical outcome. The studies vary with regard to the feedback direction (upwards/increment or downwards/decrement, marked as + or −) and its effect on different behavioural outcomes. This overview does not claim to be complete but rather constitutes the result of our extensive literature search in this field of research.

**Table 1.**
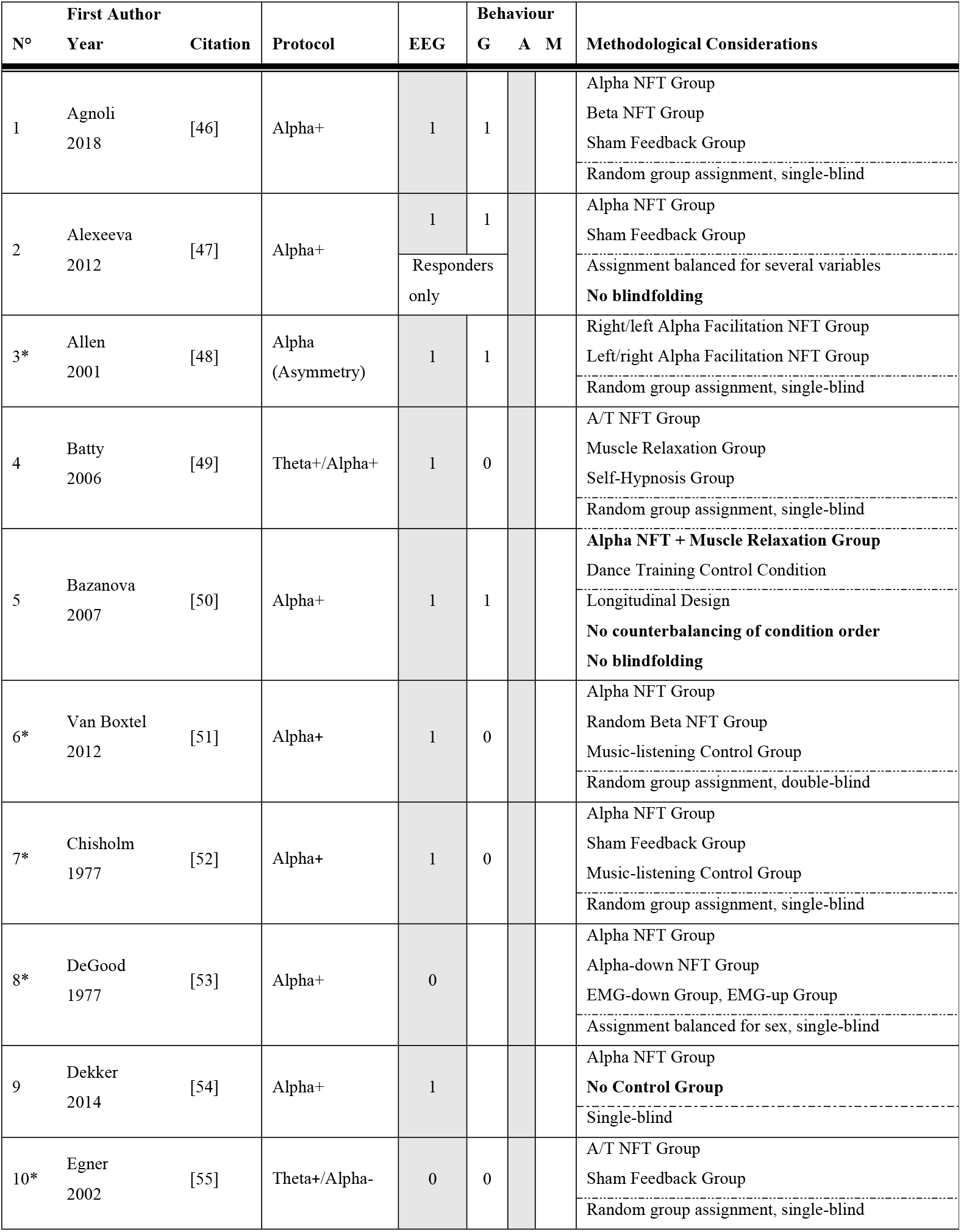
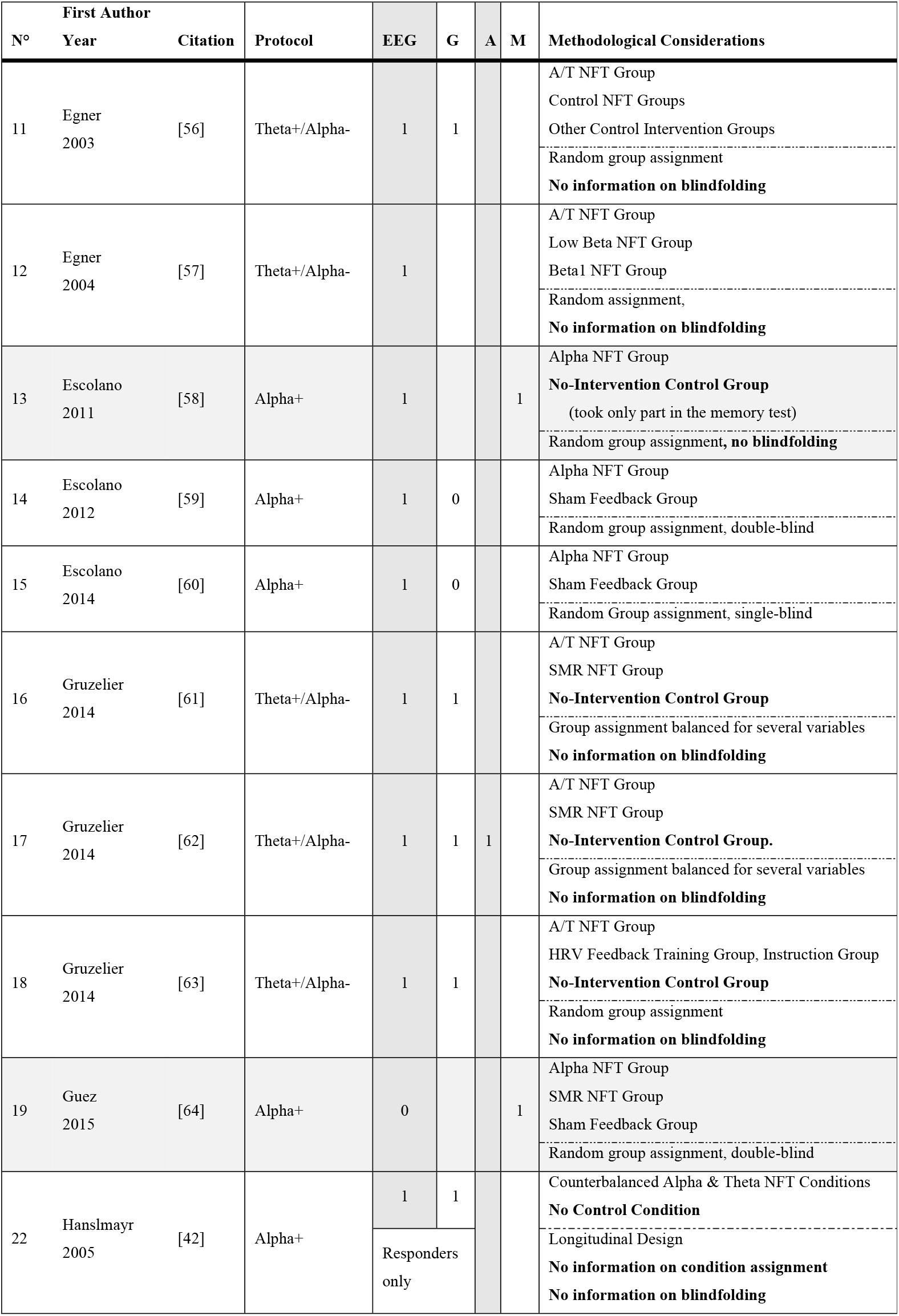
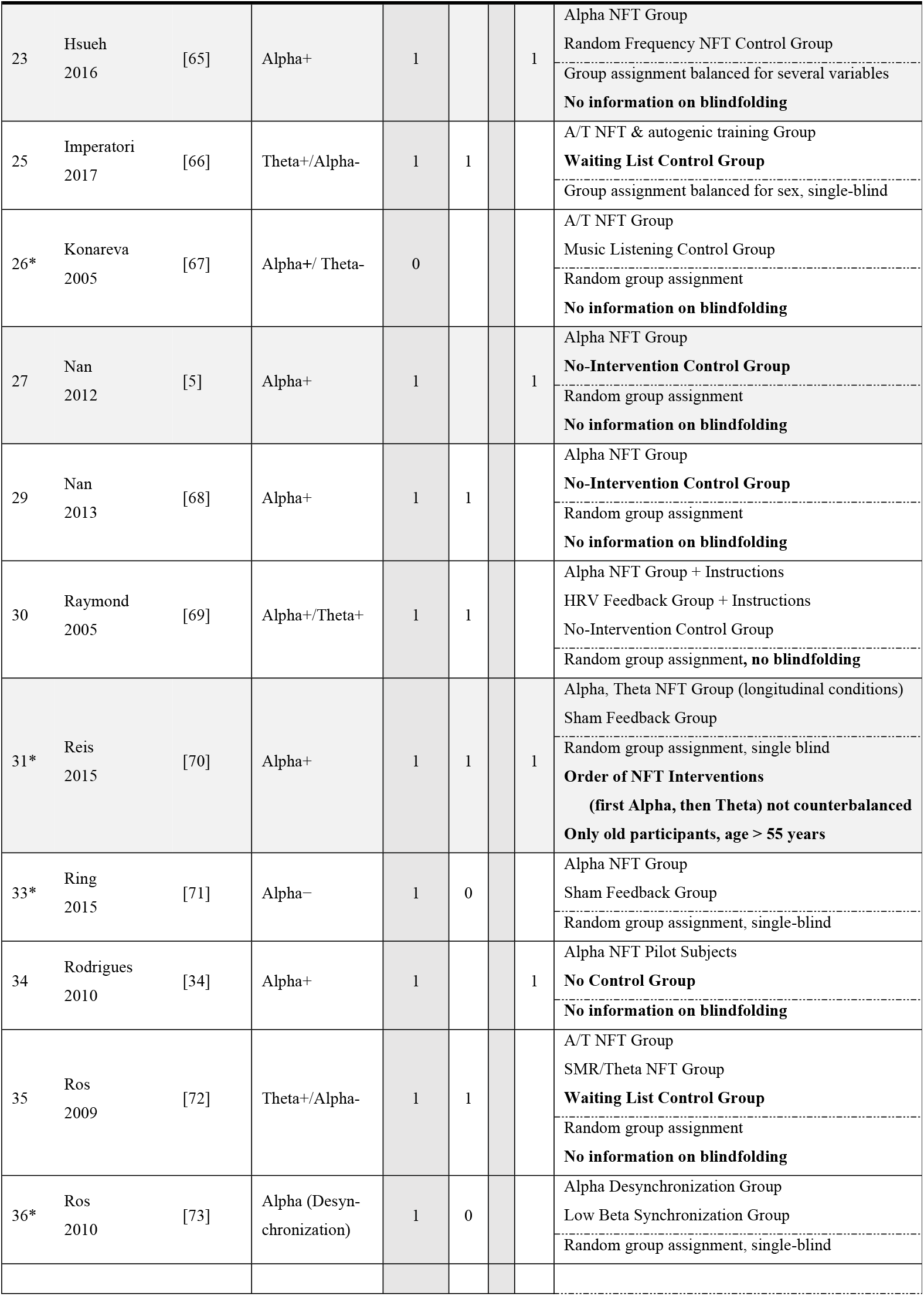
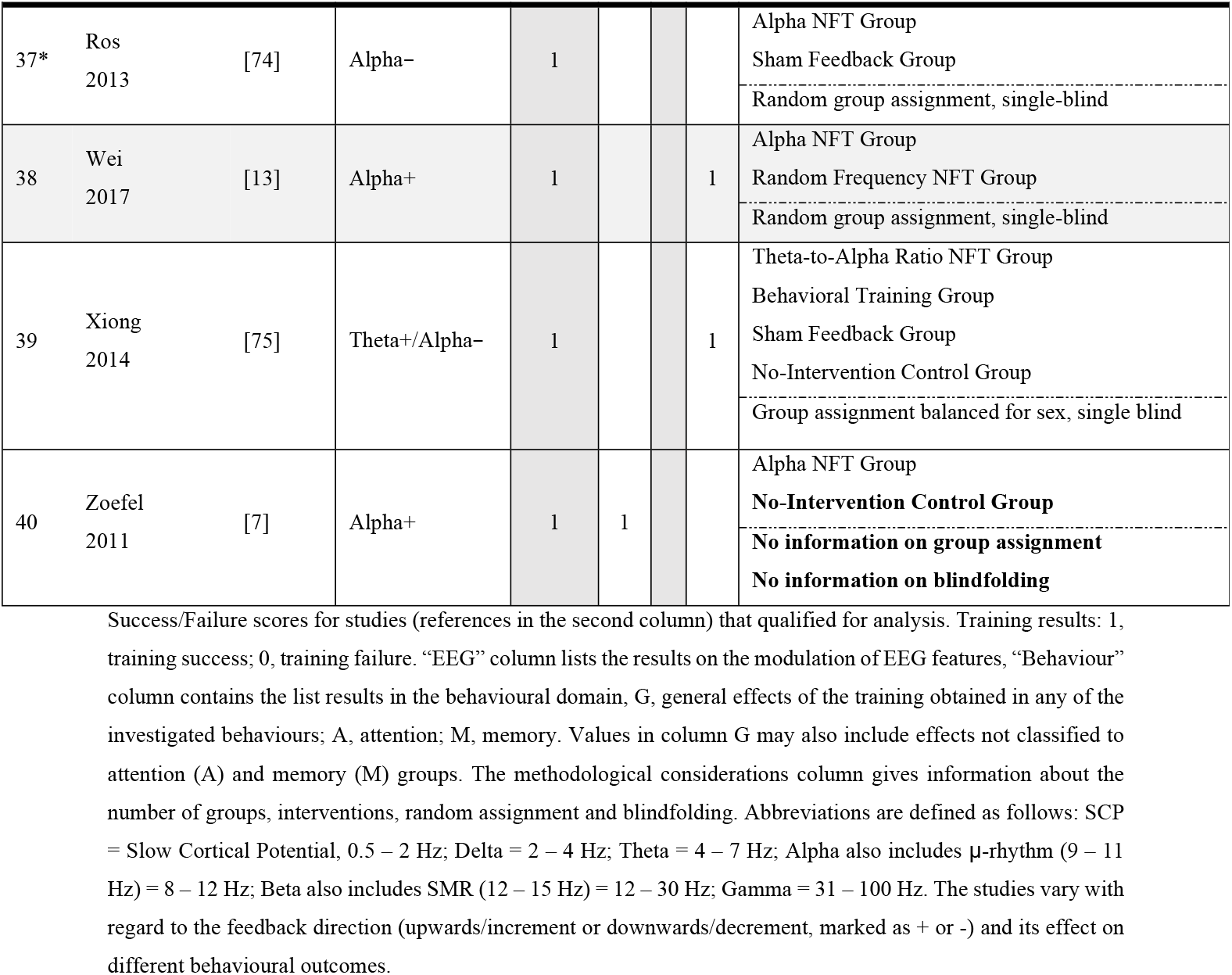
Overview of Existing Experimental Research Addressing Alpha NFT and its Effects on Behavioural Measurements

As can be observed in Table1, there is only one study using a methodologically sound experimental design with regard to a control intervention and blindfolding, which found a significant effect of alpha-up-training on memory [13]. Because of this apparent lack of evidence, this piece of research constitutes a replication study for the positive effect of alpha NFT on short-term memory performance while emphasizing the methodological aspects of the control group design and approaching the average consumer by using an easy in use and low-priced *Emotiv Epoc* EEG headset. That is why replications are needed to strengthen and better understand these initial findings.

### 1.4. The Present Study

In the past years, considerable progress was made in the development of new EEG hard- and software: today, EEG systems are available which do not require conductive gel but use saline electrodes or operate with dry electrodes instead (e.g. *Quasar*, *Neurosky or Emotiv*). Along with this simplification of physiological measurements, EEG systems are becoming increasingly user-friendly and affordable. These new user-friendly and low-priced systems do not claim to compete with state of the art high-priced EEG systems. However, they measure valid EEG signals [76,77] and have their own niche: the average consumer [78].

The study at hand combines the use of an easy to use and comparably low-priced EEG signal acquisition system (*Emotiv Epoc* EEG headset) with the question regarding the connection between alpha NFT and cognitive enhancement while adopting a methodologically sound experimental design. The following research question was formulated.

#### Does individual upper alpha NFT with an easy in use, and comparably low priced Emotiv Epoc EEG headset enhance cognitive performance in reasonably healthy participants?

In line with previous studies reporting increased individual upper alpha amplitude for NFT trainees (c.f. section 1.1 above), hypothesis H1 suggests that by means of IUA NFT, the relative IUA is enhanced. No such effect is observed in the sham feedback (SF) control group.

Referring to the results showing a significant increase in short-term memory performance after NFT (cf. section 1.2 above), hypothesis H2 suggests that alpha-NFT has a positive effect on short-term memory performance resulting in a greater increase in digit-span performance in the experimental group, compared to the control group.

In concordance with the theory of a connection between high alpha power during resting state and low alpha power during the execution of a mental task, so called event related desynchronization, hypothesis H3 is conjectured: There is an immediate positive effect of alpha-NFT on short-term memory performance. No such effect can be found in the SF control group.

## 2. Materials and Method

### 2.1 Participants

Thirty-three psychology students (26 female) were recruited at the *University of Fribourg* (Switzerland) vie E-Mail and advertisement on the campus, ranging in age from 19 to 25 years (M = 21.27 years, *SD* = 1.43 years).

After being duly informed about the protocol of the study, all participants agreed to written informed consent authorized by the ethical committee of the *University of Fribourg.* As a compensation for their participation, they earned 5 credit points on a university-intern reward system. Participants were assigned randomly to either the experimental neurofeedback training group (NFT group, n_1_ = 17, M_Age_ = 21.29, 12 female) or the control sham feedback group (SF group, n_2_ = 16, M_Age_ = 21.13, 14 female). To assess whether subjects were aware of their condition, the last experimental task was to guess which group they were assigned to. The statement ‘I was assigned to the control group’ was answered on a 7-point Likert scale (ranging from ‘I strongly disagree’ to ‘I strongly agree’). Analysis of the data showed that NFT and SF group did not differ, which indicates that participants in either group were unaware of their status (*M*_NFT_ = 2.60, *SD*_P1_ = 1.45; *M*_SF_ = 3.18, *SD*_SF_ = 0.33; t(29) = 1.18, *p* = .249).

### 2.2. Experimental Protocol

In order to control for the influence of the circadian rhythm [12], each participant was scheduled to come to the laboratory at the same time-slot on 4 consecutive days (i.e. 4 sessions, e.g. from Monday to Friday always at 10 o’clock), see Fig 1.

**Fig 1.**
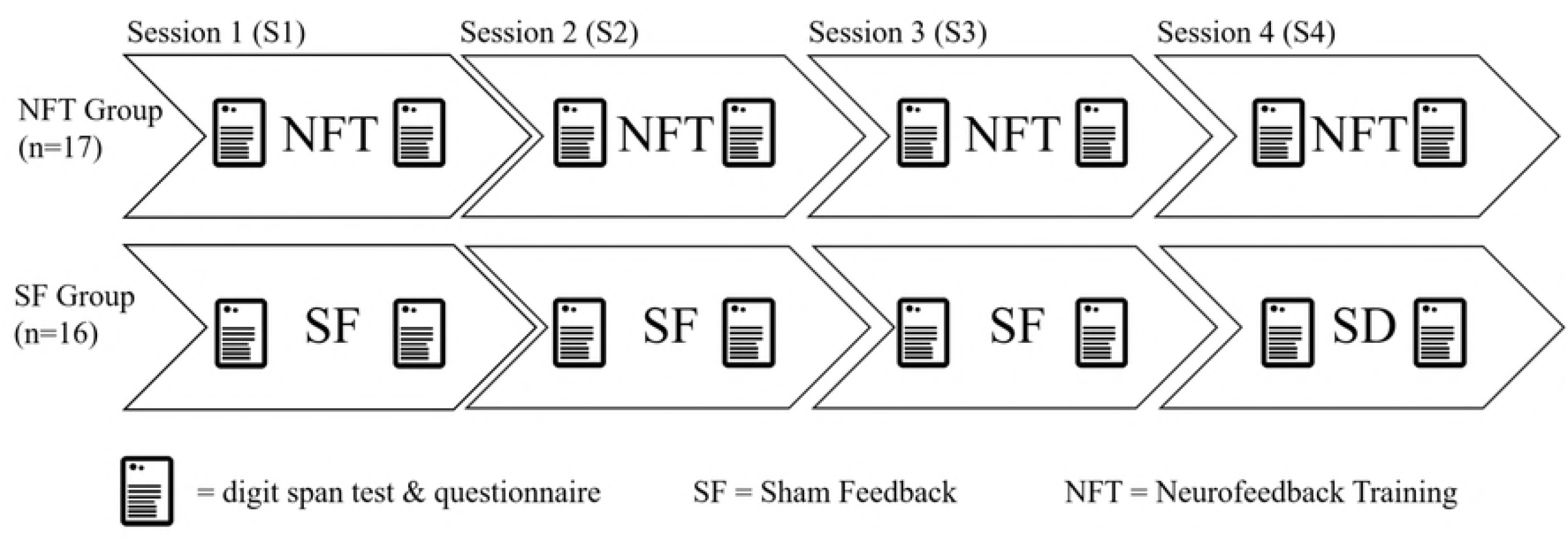
Procedure over four sessions. Experimental procedure in experimental (Neurofeedback) and control (Sham Feedback) group over four sessions on four consecutive days.

A more detailed description of the experimental session undergone from session 1 (S1) to session 4 (S4) ensues. If not indicated differently, all of the following details apply for both, the experimental group (receiving NFT) and for the control group (receiving a sham feedback intervention).

After being equipped with the EEG headset, participants filled out the MDBF mood questionnaire [79] and a series of questionnaires concerning their daily physical activities and use of substances like caffeine, alcohol and cannabis, variables that have possible implications on alpha activity [80–82]. Participants then performed a short-term memory test followed by two 2-min resting state EEG recording epochs, one with eyes closed and another with eyes open. These baseline recordings were used to assess the individual upper alpha peak for the NFT group (see next section for details).

NFT or Sham feedback (SF) started immediately after the baseline recordings and consisted of five 3-min periods with a 30 second break in-between. For S1, participants first received verbal and written information about alpha activity and were encouraged to be creative and come up with five personal strategies for the five periods of NFT (or SF). A list with five strategies (positive thinking, evoking emotions, visualizing activities, love and physiological calm) based on [5] was offered to participants who had difficulty coming up with their own ideas. Participants were asked to use only one strategy during each period, write it down during the break and to use every strategy only once over the course of the five periods. This procedure allowed to determine the most-successful alpha-enhancing strategy (one strategy, which produced the highest relative IUA for each participant). The participants were instructed to use their most successful strategy during the following sessions of S2 to S4.

At the end of each session, participants repeated the digit span test and the MDBF. For a schematic overview of the procedure during each of the sessions S1 to S4 see Fig 2.

**Fig 2.**
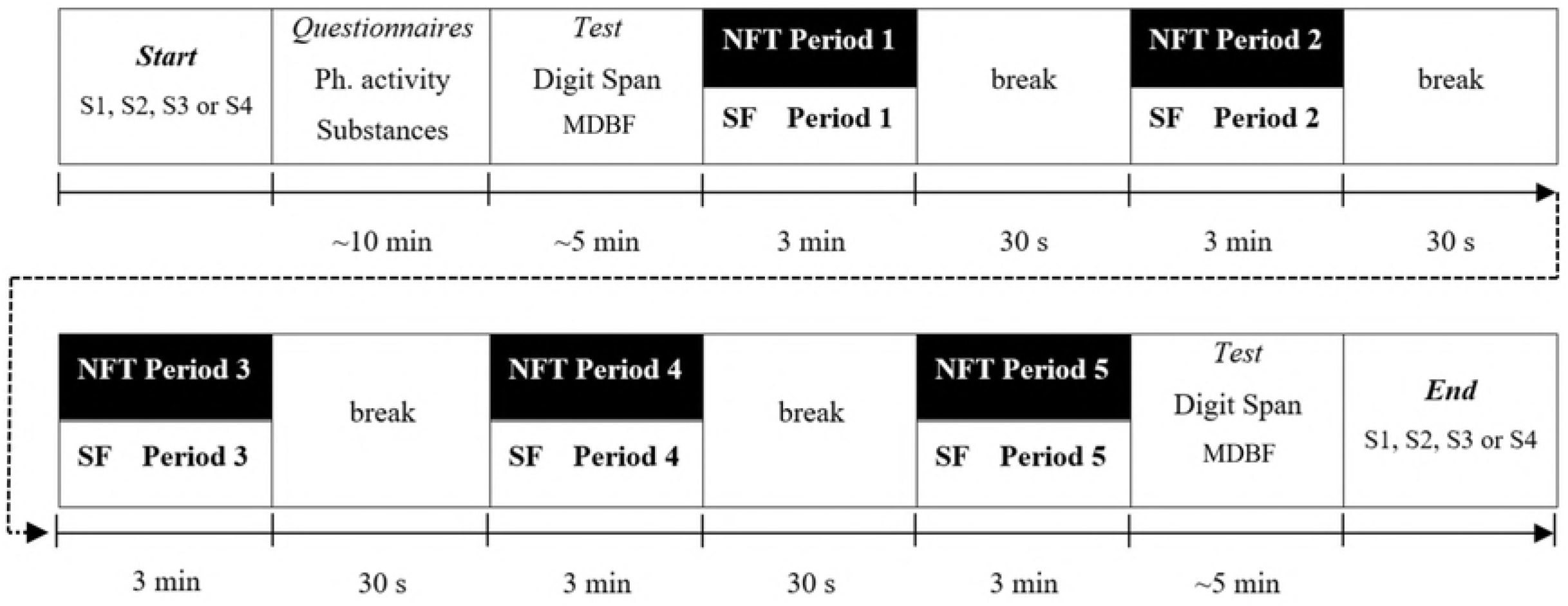
Procedure within sessions S1 to S4. Procedure during each of the sessions S1 to S4 in experimental (NFT) and control (SF) group.

### 2.3. Neurofeedback Training

Feedback sites P7, O1, O2 and P8 were chosen for their connection to visual and attentional processes (see e.g. [83,84]). Using a simple channel spectra procedure in EEGLAB (*pop_fourieeg*), each session’s baseline recordings was analysed to determine the IUA frequency band, which was then used in that session. More specifically, the individual alpha peak (IAP) between 7.5 and 12.5 Hz [12] was assessed from the eyes closed resting condition and the lower and upper border of the IUA frequency band were defined as IAP and IAP + 2, respectively. We used the *Emotiv 3D Brain Activity Map* standalone software to provide IUA feedback with a colour spectrum ranging from grey (low IUA amplitude) over green to red (high IUA amplitude). During each session’s period, participants watched their real-time IUA activity at occipital and parietal sites (P7, O1, O2, and P8) colour-coded on the surface of an animated head (see Fig 3) and were advised to produce as much red activity as possible. Participants in the experimental group performed IUA NFT always with a real-time IUA band feedback. The control group received SF by watching recordings of NFT sessions from another subject not included in this sample.

**Fig 3.**
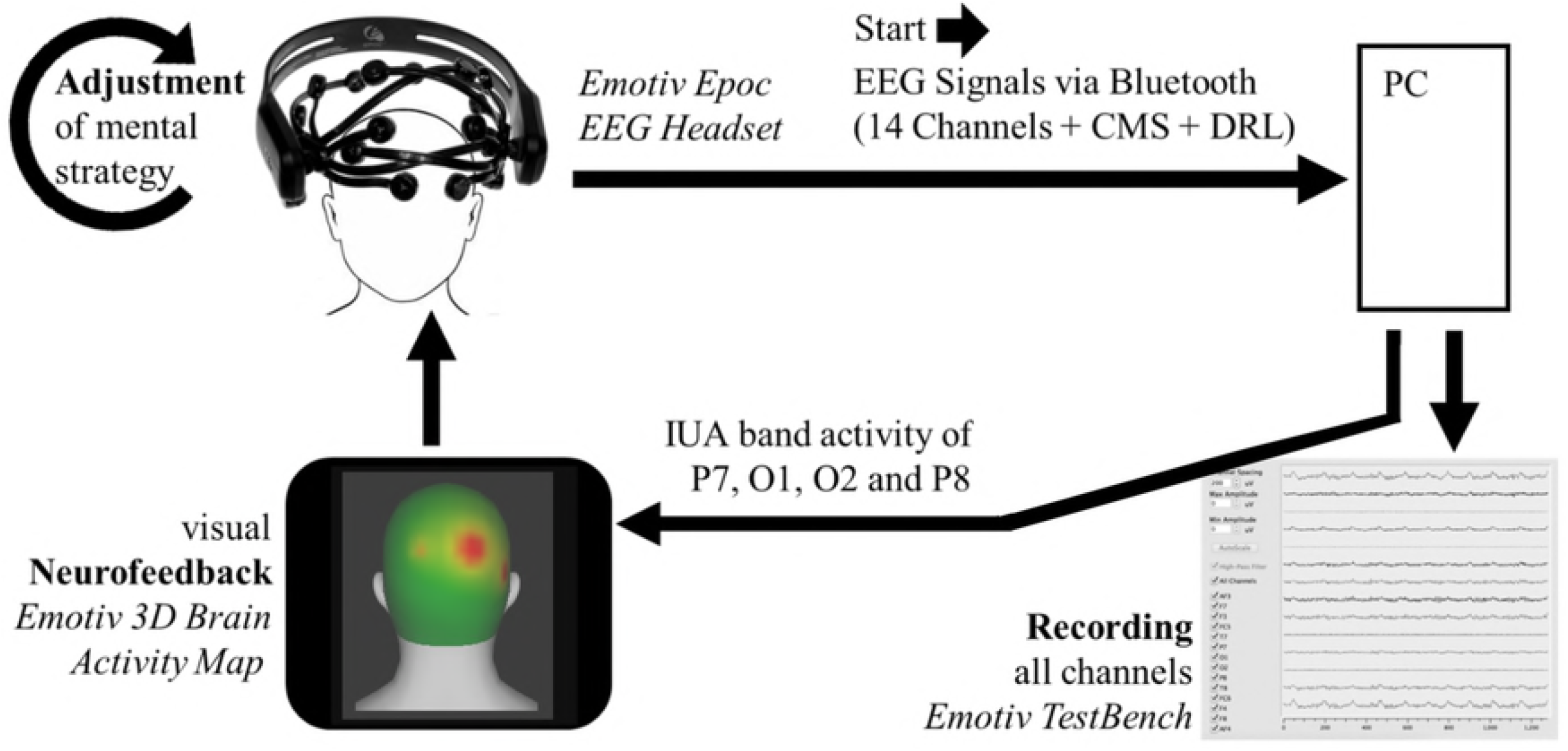
Neurofeedback loop and EEG recording.

### 2.4. EEG Recording and Processing

An *Emotiv Epoc* EEG headset was used for EEG baseline recordings and NFT. It has 14 channels (AF3, AF4, F7, F3, F4, F8, FC5, FC6, T7, T8, P7, P8, O1 and O2, international 10-20 system) and uses passive saline sensors. The device is wireless and transmits data via Bluetooth through the 2.4 GHz band, has a battery autonomy of 12 hours and uses a built-in amplifier, as well as a CMS-DRL circuit for the reduction of external electrical noise. It has a sampling rate of 128 bit/s, a bandwidth ranging from 0 to 64 Hz, automatic digital notch filters at 50 Hz and 60 Hz and the dynamic range referred to the input is 8400μV(pp). Moreover, a digital 5^th^ order Sinc filter is built-in and impedance can be measured in realtime. EEG was recorded using the software *Emotiv TestBench,* ground-reference was M1 and sampling method was by default sequential sampling.

All analyses were carried out with MATLAB and EEGLAB [85]. The data was pre-processed using the following methods: re-referencing to channel M1, automatic removal of bad epochs using the command *pop_autorej,* calculation of IC weights with the *runica* algorithm. Following [34], the relative alpha values for both, NFT and SF were calculated from the pre-processed EEG by dividing the mean amplitude of the IUA band (defined individually in the same way as in the NFT, between the IAP and IAP + 2 Hz) by the mean amplitude of the entire EEG bandwidth (Equation 1).

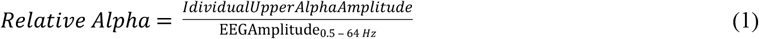

This normalization was applied to avoid variability in the absolute amplitude between trials and sessions due to changes in impedance between electrodes and scalp. This way, attenuations caused by external factors that affect all frequency bands are mitigated. Furthermore, we worked with amplitude instead of power values to prevent excessive skewing and improve the validity of the statistical analysis [6].

### 2.5. Subjective and Objective Measures

Questions about physical activities, substance intake and sleep were assessed with a self-made questionnaire. Short-term memory performance was assessed by means of a forward digit span test taken from the PEBL test battery [86]. During this test, digits appeared on the screen and participants were advised to memorize them. On first trial, three digits appeared one after another and the participant typed them into an input field in the same order as they had appeared. In case of a correct answer, a positive feedback was given and the trial was repeated with the same number of digits. If the participant succeeded again, the number of digit was increased by one. The test continued until the participant typed in a wrong answer on two consecutive trials. Two performance indicators were assessed. One is the digit span itself, defined as the highest amount of digits the participants remembered correctly. Another measure is the total correct value, representing the total number of correct answers. For example, a digit span of 9 indicates that the participant was able to remember 9 digits correctly. The total correct value in this example however can vary between 7 and 16 because participants were allowed to continue with the test if they made an error in one trial (e.g. they remembered 8 digits only once). In the statistical analysis of the present study, only total correct values are reported.

### 2.6 Statistical Analyses

All analyses were carried out with the IBM Statistical Package for Social Sciences (SPSS version 24) and R [87]. If not indicated differently, the chosen level of significance for all analyses was α = .05 (5%). Data were analysed with several mixed-measures design ANOVAs and corresponding contrast analyses using either a *polynomial* or a *simple* algorithm. Concerning the mixed design ANOVAs, *Mauchly’s Test of Sphericity* was taken into account. If *Mauchly’s Test* was not significant (*p* ≥ .20), sphericity was assumed. When *Mauchly’s Test* was significant (*p* < .20) and *Greenhouse-Geisser Epsilon* was smaller than .75, *Greenhouse-Geisser* corrected results were reported. When *Mauchly’s Test* was significant (*p* < .20) and *Greenhouse-Geisser Epsilon* was larger than .75, *Huynh-Feldt* corrected results were reported.

The general connection between alpha and digit-span was assessed with linear regressions. Additionally, for a more detailed picture, paired-samples t-tests with Bonferroni adjustment were conducted. More specifically, during each analysis (e.g. the 20×2 mixed designs ANOVA), Bonferroni correction was applied by multiplying the p-value of all associated t-tests by the number performed t-tests.

For the present study, only the change of alpha and digit-span and not their general level was of interest. Hence, all alpha measurements were standardized by subtracting the mean value of the first measurement. By applying this standardisation to experimental group and control group separately, it was assured both groups had the same initial value of alpha and digit-span, respectively. Digit-span values were not standardized because they did not differ during the first measurement (*M*_NFT_ = 8.76, *SD*_NFT_ = 1.82; *M*_SF_ = 7.94, *SD*_SF_ = 1.81; *t*(31) = 1.31, *p* = .20).

#### 2.6.1. Complementary analyses for selected extraneous variables

##### Session related changes in mood

Thirteen extraneous variables related to mood and effort were collected before and after each experimental session (see Figure 7B for details). We computed session related changes for each one of these variables and used principal component analysis (PCA) for feature extraction. The number of principal components (PCs) was chosen by interpreting the scree plot, and choosing the number of components until when diminishing returns would be obtained, guaranteed that at least 60% of variance could be explained. Each PC was then used as the response variable of a linear mixed effects model, resulting in one linear model for each PC. Each model had two random variables: subject as the random intercept and session number as the random slope. The fixed effect term was a triple interaction between period, experimental group and changes in relative alpha. Changes in relative alpha at each period were computed using an area under the curve with respect to increase (AUCi) formula described in [88], and these values were averaged for each session. Satterthwaite approximation for the degrees of freedom was used to compute *p*-values with the *lmerTest* package in the R programming environment [89].

##### Analysis of the extraneous variable pre-session relaxation

Here, we aimed at investigating if the inclusion of pre-session relaxation increases the predictive validity of the experimental group in changes for relative alpha for each session. A mixed effects model was used to predict the session average relative alpha AUCi, using each subject as the random intercept and session number as the random slope. The fixed effect term was the moderation between experimental group and pre-session relaxation. The moderation was introduced to understand if pre-session relaxation increases in alpha would be specific to one of the experimental groups. If the interaction term was not significant, we would test the additive model. For the latter, pre-session alpha would be tested as a suppressor variable. Satterthwaite approximation for the degrees of freedom was used to compute p-values with the *lmerTest* package in the R programming environment.

## 3. Results

### 3.1 Alpha

Regarding the temporal development of individual upper alpha (Fig 4), visual inspection of the data indicates that both groups increased in alpha. However, this impression was not confirmed by the results of a mixed measures design 20*2, TIME_α_*GROUP ANOVA (*F*(5.24, 162.36) = 1.79, *p* = .114, η_p_^2^ = .06). Moreover, neither the interaction TIME_α_*GROUP, *F*(5.24, 162.36) = 0.58, *p* = .363, η_p_^2^ = .02 nor the effect of GROUP were significant, *F*(1, 31) = 0.10, *p* = .757, η_p_^2^ = .00

**Fig 4.**
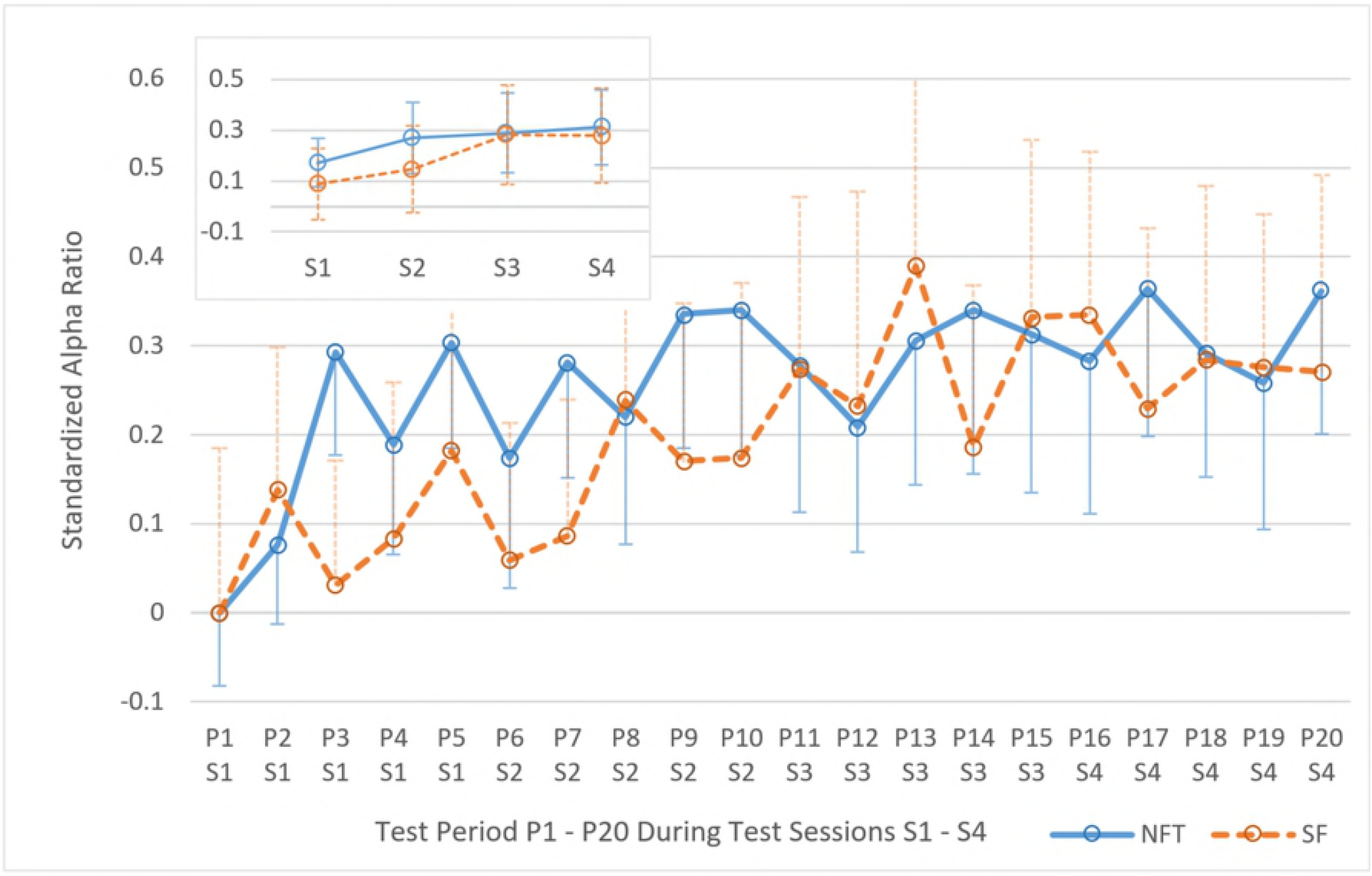
Temporal Development of Individual Upper Alpha. Temporal development of relative individual upper alpha over twenty 3-min periods of Neurofeedback Training (NFT, blue line) and sham feedback (SF, orange dashed line) during the four test sessions on four consecutive days. Relative alpha was obtained by dividing the average amplitude of the individual upper alpha band (around 10 to 13.5 Hz) by the average amplitude of the entire EEG band (i.e. 0.5 to 64 Hz). Moreover, relative alpha values was standardized with the first measurement (i.e. period 1). Error bars indicate standard error of the mean (*SEM*).

However, Fig 4 shows stronger increase between P1 and P20 for the NFT group compared to the SF group. *T*-tests with Bonferroni adjustment showed significant differences from P1 to P20 for the experimental group (*M*_P1_ = 0.00, *SD*_P1_ = 0.34; *M*_P20_ = 0.36, *SD*_P20_ = 0.66), *t*(16) = 2.63, *p* = .018, but not for the control group (*M*_P1_ = 0.00, *SD*_P1_ = 0.74; *M*_P20_ = 0.27, *SD*_P20_ = 0.88), *t*(15) = 1.33, *p* = .204. Applying *contrast* analysis (simple), the first significant amplitude difference in the SF group appeared between P1 and P11, *F*(1, 15) = 5.08, *p* = .04, *η*_p_^2^ = .25. The NFT group showed its first significant differences already between measurements P1 and P3, *F*(1, 16) = 12.67, *p* = .003, *η*_p_^2^ = .44. Contrast analyses indicate hence a faster increase of relative alpha in the NFT group.

Moreover, in the experimental group t-tests with Bonferroni adjustment showed significant improvements from P1 (*M*_P1_ = 0.00, *SD*_P1_ = 0.34) to P3 (*M*_P3_ = 0.29, *SD*_P3_ = 0.48), *t*(16) = 3.56, *p* = .006, and from P1 to P5 (*M*_P5_ = 0.30, *SD*_P5_ = 0.49), *t*(16) = 3.69, *p* = .004. The significant increase in alpha from P1 to P3 and from P1 to P5 could not be observed in the control group (*M*_P1_ = 0.00, *SD*_P1_ = 0.74; *M*_P3_ = 0.03, *SD*_P3_ = 0.56; *M*_P5_ = 0.18, *SD*_P5_ = 0.73), *t*(15) = 0.28, *p* = 1; *t*(15) = 1.08, *p* = .594, indicating a faster increase in relative alpha in the NFT group as well.

Interestingly, regardless of group and on an exploratory note, a significant positive correlation between the unstandardized initial relative alpha (P1) and the alpha improvement during the course of the experiment (P20 minus P1) was observed, *r*(31) = .44, *p* = .011. This finding was supported when the same analysis was performed on the level of test days (sessions). The initial relative alpha during the first Period (S1) correlated with the mean improvement over the course of the experiment (S4), *r*(31) = .52, *p* = .002. In other words, participants who exhibited a high relative alpha in the beginning of the experiment had a higher gain in alpha during the training compared to participants who started with a low alpha level.

The findings examined so far are partially in concordance with hypothesis H1, stating a positive effect of NFT on relative IUA. The IUA increase in the experimental group is observed earlier and the difference between P1 and P20 shows significance only in the NFT group. Interestingly and contrary to our expectation, alpha enhancement could be observed in the control group as well, when contrast analyses are taken into consideration.

### 3.2. Neurofeedback Training and Short-Term Memory Performance

Regarding the temporal development of STM performance, a significant main effect of TIME_stm_ was observed, *F*(7, 217) = 4.90, *p* < .001, η_p_^2^ = .14, but the interaction TIME_stm_*GROUP did not reach significance level, *F*(7, 217) = 1.24, *p* = .280, η_p_^2^ = .04. No effect of factor GROUP was observed, *F*(1, 31) < 0.01, *p* = .963, *η*_p_^2^ < .01. Contrasts showed a linear trend of TIME_stm_ with *F*(1, 31) = 6.36, *p* = .017, *η*_p_^2^ = .17. No linear trend of the interaction TIME_stm_*GROUP was observed *F*(1, 31) = 0.02, *p* = .887, *η*_p_^2^ < .01.

Paired-samples t-tests with Bonferroni adjustment revealed no significant differences between first and last measurement of STM in the experimental group (*M_TI_* = 8.77, *SD_TI_* = 1.82; *M_T8_* = 10.06, *SD_T9_* = 2.49), *t*(16) = 1.85, *p* = .083, but for the control group (*M_TI_* = 7.94, *SD_TI_* = 1.81; *M_T8_* = 9.69, *SD_T8_* = 1.96), *t*(15) = 2.78, *p* = .014. All results examined in this section contradict hypothesis H2 postulating a general effect of NFT on STM.

To evaluate the immediate effect of NFT on STM performance, a 2*2 mixed-model ANOVA with the within factor PRE/POST and the between-groups factor GROUP was conducted. Factor PRE/POST had two levels: averaged digit span performance values conducted *before* the intervention vs. averaged digit span performance values conducted *after* the intervention (see Fig 7). Participants did not improve in STM performance during NFT/SF, *F*(1, 31) = 2.98, *p* = .094, *η*_p_^2^ = .09. Test of within subjects effects did not reveal an interaction effect, *F*(1, 31) = 1.26, *p* = .271, *η*_p_^2^ = .04. No effect of GROUP was observed, *F*(1, 31) = 0.02, *p* = .907, *η*_p_^2^ < .01.

**Fig 5.**
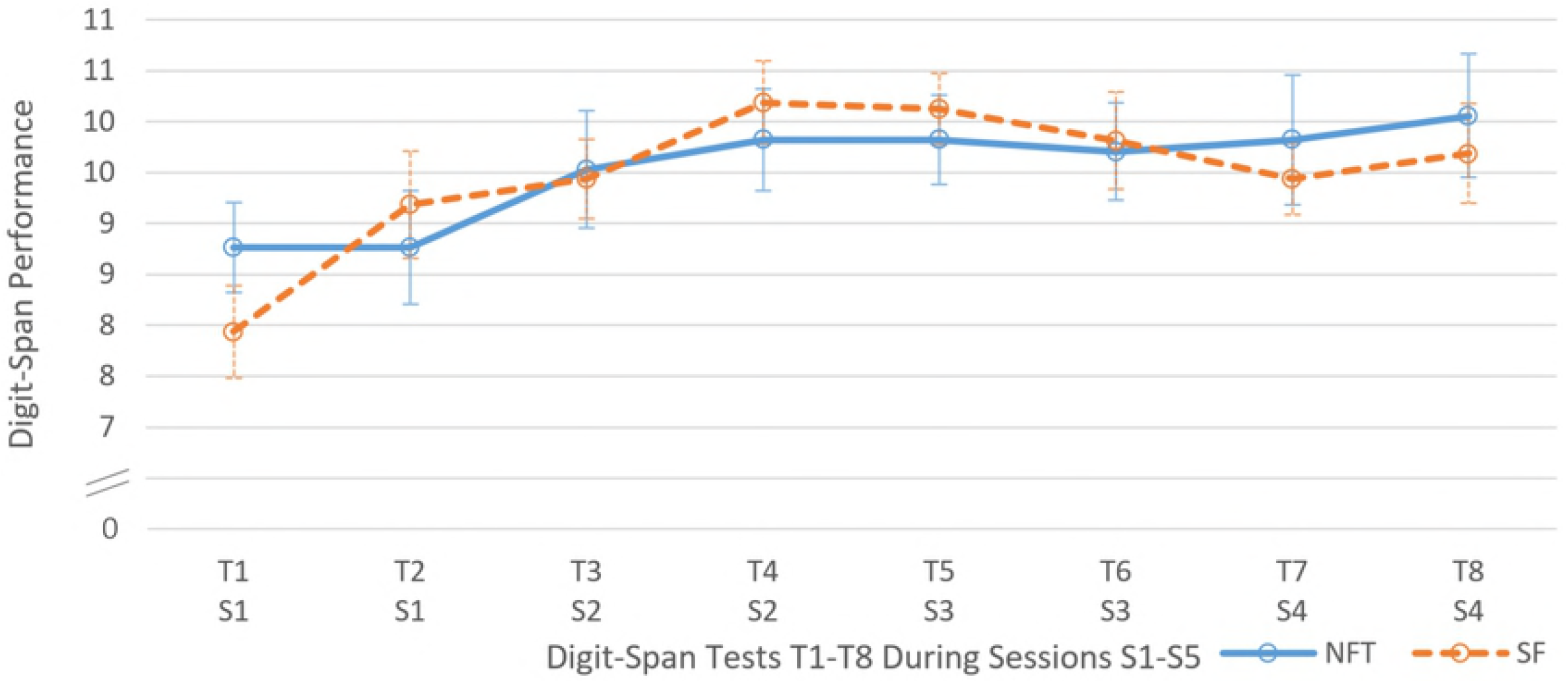
Temporal development of STM performance. Temporal development of short-term memory (digit-span) performance over 8 tests (T1 to T8) in a neurofeedback (NFT, blue line) and a sham feedback (SF, orange dashed line) group. Subjects participated in four consecutive test days (S1-S4) containing two tests each. Uneven test numbers (T1, T3, T5 and T7) were conducted before the intervention (NFT or SF), even test number (T2, T4, T6 and T8) were conducted after the intervention. Error bars indicate *SEM*.

**Fig 6.**
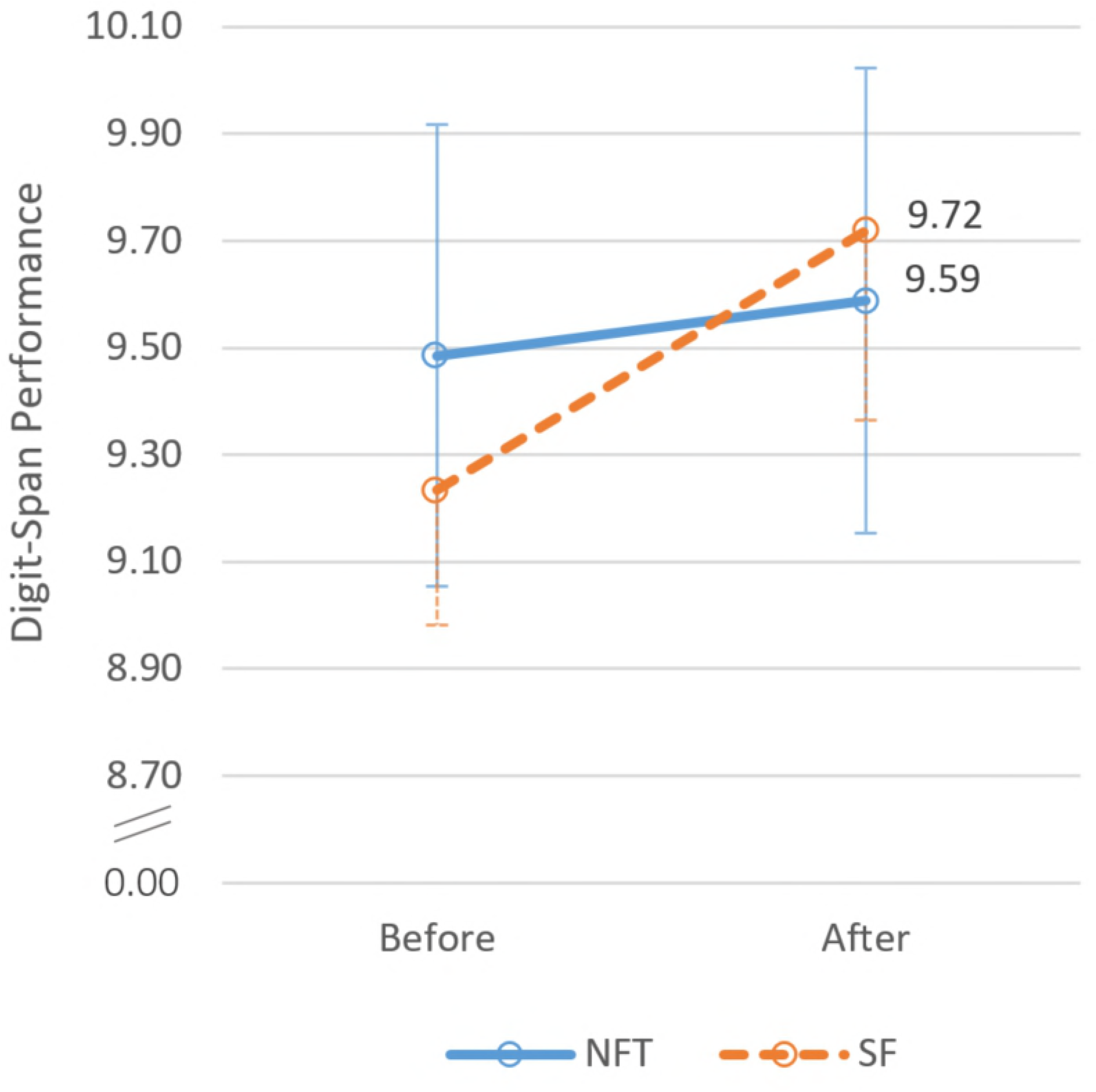
STM performance before vs. after the intervention. Short-term memory performance, measured by digit-span tests *before* and *after* neurofeedback training (NFT, blue line) and sham feedback (SF, orange dashed line). Error bars indicate *SEM*.

**Fig 7.**
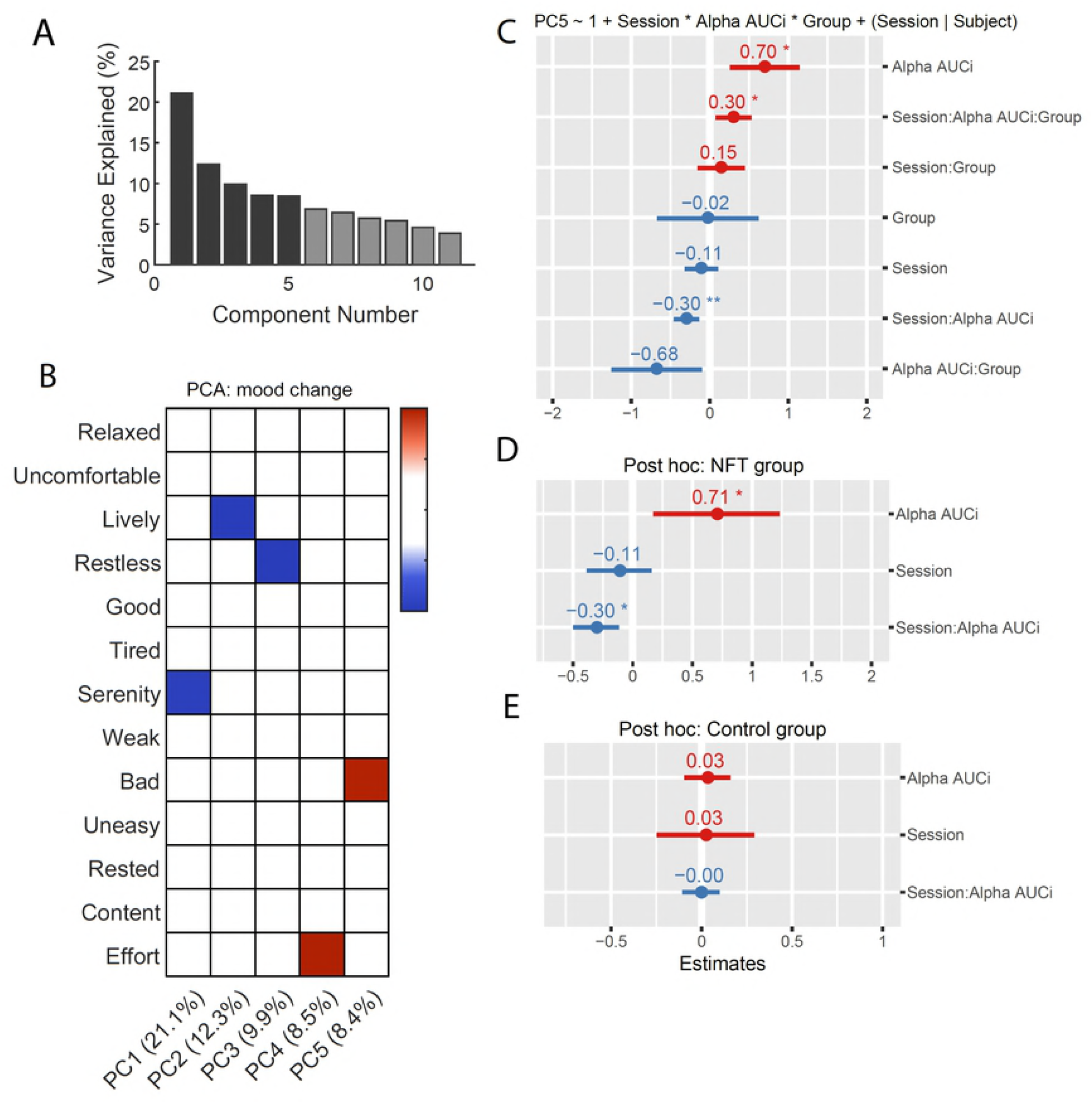
Effects of NFT training on mood change. A) Percentage of variance explained for each PC. The 5 PCs used result in a total of 60.2% of variance explained (dark-shaded bars). B) Loading matrix for each PC after Varimax rotation. Loadings smaller than 0.4 are not shown. C) Linear mixed effect model for PC5 as the response variable and the triple interaction between session, relative alpha AUCi and experimental group as predictors. D) Simple effects model for the NFT group. E) Simple effects model for the control group. For panels C), D) and E) coefficient estimates and standard errors (SE) depicted as dot and line respectively. Red and blue colors represent positive and negative coefficient estimates, respectively. Significance levels: ** *p* < .01, * *p* < .05. Significance levels are presented for uncorrected *p*-values. When Bonferroni correction is applied, to control for Type I errors due to the comparing models for five PCs, the triple interaction term in panel C is no longer significant (*p*corrected = 0.152). For panels D and E, when applied, Bonferroni correction controls for two comparisons made in the simple main effects analysis resulting in a significant interaction effect ‘Session x relative alpha’ AUCi (*p*_corrected_ = 0.028).

These findings do not support hypothesis H3 postulating an immediate positive effect of NFT on STM.

### 3.3. Alpha and Short-Term Memory

To assess the connection between alpha and STM, a multiple regression analysis was conducted. The dependent variable was STM improvement defined as performance delta-value of test 8 (T8) minus test 1 (T1). Relative alpha values were averaged over sessions (see upper-left corner Fig 4) and served as predictors. Explained variance R^2^ was 0.10 and the corresponding ANOVA was not significant, *F*(4, 32) = 0.79, *p* = .540. This finding contradicts hypothesis H2, assuming a general connection between alpha and STM performance

### 3.4. Analyses for selected extraneous variables

#### 3.4.1. Session related changes in mood

In order to infer how NFT affected mood changes during the experiment, we calculated the differences in mood from the beginning to the end of each session. We also calculated total amount of change in relative alpha at each period using an area under the curve with respect to increase (AUCi) formula described in [88]. The 12 mood change variables, and one variable representing effort, were compressed using principle component analysis (PCA) into 5 principal components (PCs) explaining 60.2% of the variability in the original variables (See Fig 7A). *Varimax* rotation was used to facilitate interpretation of each PC, resulting in the loading matrix in Fig 7B. Each PC was used as the response variable in a linear mixed effects model (see Figure 7C for the model with PC5 as the response variable) with each subject as the random intercept, session number as the random slope and a triple interaction between period, experimental group and relative alpha AUCi. The only PC with a significant (*p* = 0.030, however when applying Bonferroni correction for 5 comparisons *p*_corrected_ = 0.152) triple interaction predictor was PC5, a component that loads positively on the variable changes in the bad mood (positive values of PC5 represent an increase in bad mood). This model had a total explanatory power (conditional R^2^) of 48.06%. The triple interaction between session, relative alpha AUCi and experimental group (see Figure 7C; β = 0.30, SE = 0.14, 95% CI [0.035, 0.57], t(105) = 2.20, *p* = .030, *p*_corrected_ = 0.152) could be considered a small effect (std. β = 0.33, std. SE = 0.15). Simple main effects for each experimental group were analyzed in order to evaluate whether the interaction of session and relative alpha AUCi was significant only for the NFT group (after correcting for these two comparisons). For this group, the interaction effect between Session and relative alpha AUCi was significant (β = −0.30, SE = 0.12, 95% CI [−0.53, −0.070], *t*(52) = −2.56, *p* = .014, *p*_corrected_ = 0.028) and could be considered as small (std. β = −0. 25, std. SE = 0.097). Since no effects were found for the control group (*p* = 0.997, *p*_corrected_ = 1), these results suggest that in the NFT group, learning to progressively increase the relative alpha band lead to small but significant reductions in bad mood.

#### 3.4.2 Analysis of the extraneous variable pre-session relaxation

One hypothesis to explain the similar alpha production between the NFT and control group is, that participants in the control group, although not receiving real feedback, were also trying relaxation strategies (since this was one of the cognitive strategies recommended to participants). Therefore, we decided to investigate if the experimental group variable would be capable to predict higher relative alpha AUCi values in the NFT group by accounting for the moderation between experimental group and pre-session relaxation. Fitting the model described in Figure 8 to the data, the effect of experimental Group was significant (β = −1.89, SE = 0.72, 95% CI [−3.30, −0.47], *t*(94) = −2.63, *p* < .010) and could be considered small (std. β = −0.20, std. SE = 0.17). The negative value of the estimated coefficient points to a lower overall relative alpha AUCi for the control Group. The effect of the interaction between level of relaxation at the beginning of the session and experimental Group (β = 0.47, SE = 0.19, 95% CI [0.088, 0.84], *t*(99) = 2.45, *p* = .016) and could be considered small (std. β = 0.44, std. SE = 0.18). This indicates either the NFT or the control group could show an effect of level of relaxation on the production of relative alpha. A post hoc analysis revealed that this is not true for the NFT group: The effect of being relaxed was not significant (β = −0.19, SE = 0.11, *t*(37) = −1.63, *p* = .112). For the control group there was a trend for being relaxed leading to more relative alpha increases (β = 0.28, SE = 0.15, 95% CI [−0.029, 0.58], *t*(53) = 1.87, *p* = 0.068) and the effect could be considered small (std. β = 0.23, std. SE = 0.12). These results suggest that if the level of relaxation before each Session is taken into consideration, then it is possible to observe an overall effect of NFT on the production of relative alpha. It also suggests that for the control group, being relaxed might have been what lead to increases in relative alpha.

**Fig 8.**
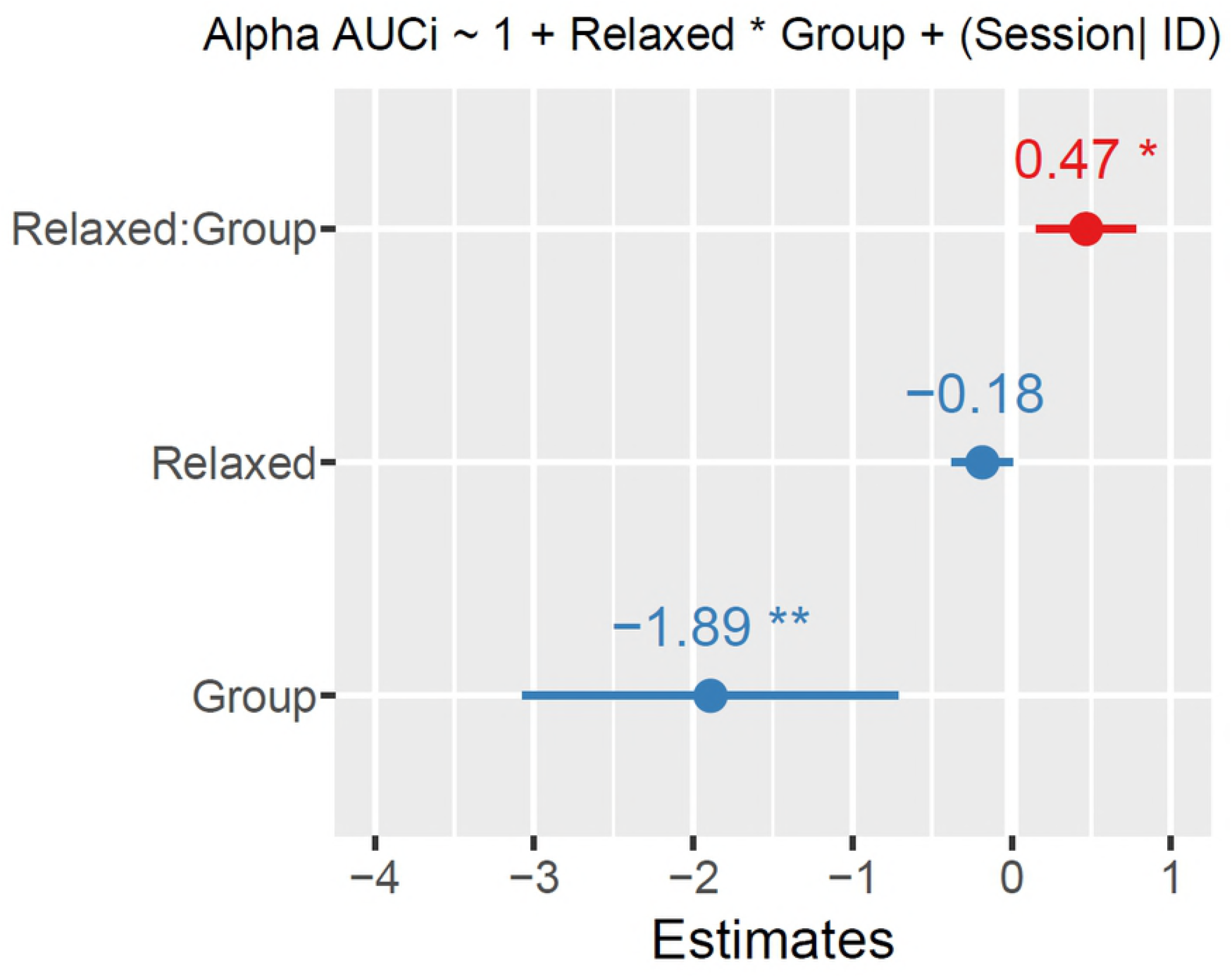
Linear mixed effect model for relative alpha AUCi as the response variable and the interaction between how relaxed participants were at the beginning of the Session and experimental group as predictors. Coefficient estimates and standard errors (SE) depicted as dot and line respectively. Red and blue colors represent positive and negative coefficient estimates, respectively. Significance levels: ** *p* < .01, * *p* < .05.

## 4. Discussion

The present study investigated the connection between IUA alpha NFT, relative alpha and short-term memory performance using a commercially available BCI device (*Emotiv Epoc*) in a single-blind design. In line with previous results [5], an enhancing effect of the training on the relative alpha and on short-term memory performance (*digit-span Task*) was expected.

Our analyses showed a significant improvement in relative alpha in the neurofeedback group between period 1 and period 20 which could not be observed in the sham feedback group. Moreover, contrasts showed that the increase in alpha was obtained much earlier (period 3) for participants who saw their real-time brain activity compared to participants who followed a sham-feedback intervention (period 11). Additionally, we also observed that if the level of relaxation before each Session is taken into account, it is possible to observe a clear effect of NFT on the production of relative alpha. All in all, the results regarding relative alpha indicate that up-training alpha with a real-time NFT procedure facilitates the process of enhancing alpha activity. Thus, the results of the present study were in accordance with hypothesis H1.

Furthermore, we hypothesised the control group would not show an enhancement in alpha activity. Interestingly though, contrast analyses showed the opposite was true and mixed measure longitudinal analysis indicated that the slope for both groups were equal. One possible explanation for this finding is that the control group was given feedback during the first session. Although sham feedback was used for this group, by the end of this session they were informed of the most successful mental strategies for alpha upregulation. This would imply that it is possible to infer appropriate mental strategies within one session and that coherent visual feedback might not be necessary for ensuing sessions to upregulate alpha to a certain degree, provided the adequate mental strategy is used. This interpretations seems to be in line with studies showing an alpha enhancing effect of certain types of meditation (e.g. [48–50]). A replication study with an additional control group that would not be informed about which strategies are generally linked to alpha enhancement might address this question. Another possible explanation resides in the framework of socio-physiological processes. Alpha is enhanced by being in a calm and resting state and in the course of the current study, participants became more and more familiar with the experimental environment, as well as with the experimenter. It is likely that the participants became more and more relaxed, comfortable and calm during the later sessions of NFT/SF, which might have led to the observed enhanced level of alpha in the control group. This interpretation finds some support in the analysis of extraneous variables, which suggests that being relaxed is an important factor for increases in relative alpha. In line with [93] it could be important to collect data from a control group advised with an inverse NFT paradigm.

No general connection between alpha and STM performance was observed in this study and thus, no evidence supported hypothesis H2. There was indeed a significant effect of time throughout the eight digit-span tests but no main or interaction effects due to NFT. There was also no immediate positive effect of NFT on STM performance. All in all, these findings contradicted our hypotheses and several explanations can be put forward.

For one, the quick improvement in the digit-span task observed in most participants is most likely due to practice and it seemed to be stronger than the more subtle positive impact of higher alpha amplitude. Electrode placement could also be responsible for the lack of effect of NFT on STM performance. Feedback signals were acquired from P7, O1, O2 and P8 because the occipital cortex is involved in every visual process and parietal sites are connected to attention (e.g. [45]). It is possible though, that the choice of electrodes influenced or impaired the effect of the NFT on STM performance. Many authors use electrode sites like Cz, Pz, Fz and C3 which differ from the sites used during the current study (see e.g. [9,13,51,52]). However parietal and occipital electrode sites are used commonly for IUA NFT as well (e.g. [10,36,53]). Finally, the possibility of the absence of an effect of IUA NFT on STM performance should be considered. Previous studies where this effect was reported have used no-intervention or waiting-list control groups (e.g. [5,10]), which deserve critical attention. Neither have they ruled out the expectancy effect on the side of the participants (placebo or Hawthorne effect, [94]), nor did such designs compensate for a potential experimenter bias. In the few studies using randomized sham feedback control groups, no significant results indicating an effect of alpha feedback on STM performance could be reported [60,95]. All these observations are in accordance with the review of Rogala [45], which excluded many alpha NFT studies for methodological considerations and could rate only one study [70] as success in regard of the effect of NFT on memory (i.e. *digit-span task*).

Nevertheless, some limitations in this study need to be pointed out which could have obfuscated this effect. Although it is used in various clinical test batteries and generally is considered a useful indicator for cognitive performance [96], the digit-span task used in this study showed rather low intra-individual variation and strong learning effects in the repeated measures design. Therefore, a different measure for cognitive performance (e.g. Mental Rotation Task, N-Back or a Trail Making Test) might have led to larger variances without concealing a potential NFT effect by learning.

It is also possible, that the conditioning paradigm was not efficient enough due to using a simple colour-code as feedback signal. Other authors worked with very specific reward symbols and sounds (e.g. beeps, counters, pleasant sounds or even applause, [25,48,50]). Future implementations should guarantee that NFT is done in an immersive environment with clear and intuitive rewards. This is particularly important in the perspective of NFT with commercially available devices since they most probably will be used outside controlled laboratory settings.

Finally, the study also has some limiting aspects regarding how mental strategies were employed to enhance individual upper alpha activity. Participants were asked to maintain the same strategy after the first training session. Interestingly, the corresponding gain in alpha activity on the first day was very high compared not only to the first measurement of NFT but also compared to the rest of the training course. In period 5 (P5), participants in the NFT group reached nearly the same relative alpha (*M*_IUA,P5_ = 0.30, *SD*_IUA,P5_ = 0.49) as in the last period, P20 (*M*_IUA,P20_ = 0.36, *SD*_IUA,P20_ = 0.66). Perhaps, choosing one successful strategy for alpha enhancement is not as effective as advising the participants “to be guided by the feedback process” itself [56] or to interchange between more than one strategies in order to avoid fatigue or, the effect of alpha NFT plateaus in general after a certain amount of training like Dekker states [54].

One of the main motivations for this study was to assess if a commercially available EEG device could offer the necessary means to achieve EEG self-regulation and, in turn, reap its benefits in cognitive improvement. We think this is an important question to answer given the promising benefits of NFT on mental health and well-being, its non-invasiveness, and the growing affordability of commercial EEG devices; NFT is no longer just important in the clinical setting, but also in real-life settings. Although our hypotheses for cognitive improvement were not verified, we observed promising results in our experimental group concerning alpha upregulation: this group achieved significant alpha increases faster and these increases were associated with decreases in negative mood. We expect this study to be a stepping stone in larger collection of works that aim for ecological validity and sound experimental design. Ultimately, it will be possible to collect data from a large number of participants following NFT at their homes, workplaces or any place of their choosing so, it is urgent to start defining appropriate protocols. While the NFT implementation of the present study might not be suitable for the daily use, NFT harbours great potential, especially, in our opinion, when combined with gamification strategies. This way, the enhancing effect of NFT could be combined with the immanent beneficial effect of computer games (see e.g. [57]) and immersive virtual environments. Future research should not only focus on theoretical aspects of the working mechanism behind NFT, but on the development of engaging NFT implementations with practical relevance.

